# Multiplexed single-cell RNA-seq via transient barcoding for drug screening

**DOI:** 10.1101/359851

**Authors:** Dongju Shin, Wookjae Lee, Ji Hyun Lee, Duhee Bang

## Abstract

To simultaneously analyze multiple samples of various conditions with scRNA-seq, we developed a universal sample barcoding method through transient transfection of SBOs. A 48-plex drug treatment experiment of pooled samples analyzed by a single run of Drop-Seq revealed a unique transcriptome response for each drug and target-specific gene expression signatures at the single-cell level. Our cost-effective method is widely applicable for single-cell profiling of multiple experimental conditions.

---

Unlike conventional bulk measurements, single-cell RNA-sequencing (scRNA-seq) helps analyze transcriptomes of individual cells^1–3^, and has shed light on variations in cell populations, such as tumor heterogeneity. Platforms such as Drop-Seq^4^, inDrop^5^, and 10X Genomics Chromium^6^ provide high-throughput single-cell information over thousands of cells. Although it is necessary to carry out scRNA-seq on samples of various conditions or of many patients, the previous scRNA-seq method about multiple samples is still challenging because of a labor-intensive process and high sample-preparation costs. To overcome these limitations, we designed a multiplexed scRNA-seq method that involved transient transfection of short barcoding oligos (SBOs) to label samples from various experimental conditions.

SBO, a single-stranded oligodeoxynucleotides, consists of a sample barcode and a poly-A sequence **(Supplementary Fig. 1a)**. Transient transfection of SBO allows a sample to be labeled with a unique barcode. The barcoded samples of various conditions are pooled and simultaneously processed for scRNA-seq (Fig. 1a). The poly-A sequence in the SBOs ensures that the mRNAs and SBOs are captured and reverse-transcribed together during the scRNA-seq process **(Supplementary Fig. 1b)**. Computational analysis of digital count matrices of SBOs allows us to demultiplex and determine sample origins. This universal barcoding method, based on simple transfection enabled sample multiplexing, identified multiplets and negatives, and reduced the preparation cost per sample.

**Figure 1.**
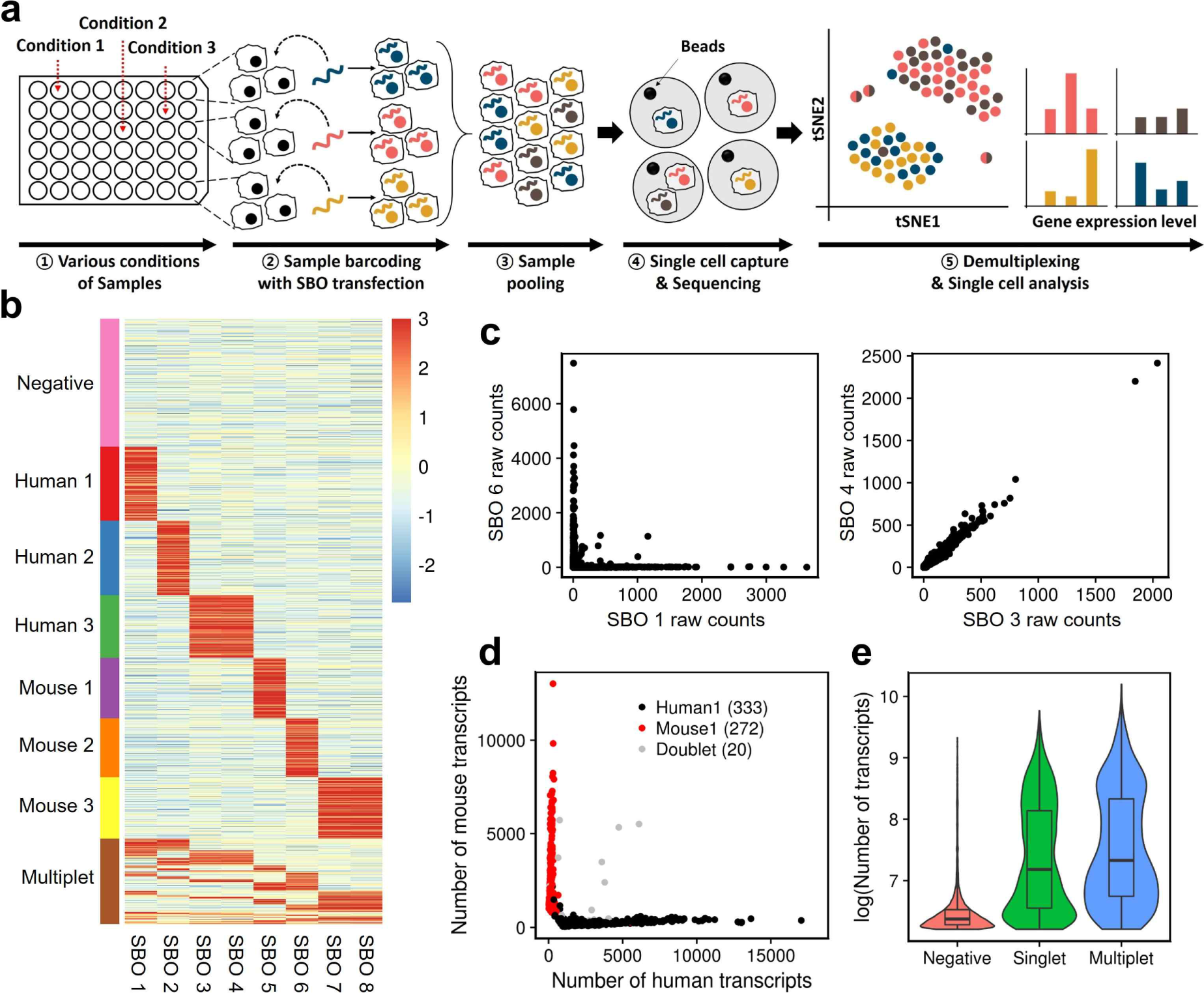
Scheme and validation of transient barcoding method. **(a)** Scheme of multiplexed scRNA-seq by transient barcoding method using SBOs. (1) Various conditions of samples are prepared. (2) Each sample is transfected with SBO containing a unique sample barcode. (3) Barcoded cells are pooled altogether and processed for scRNA-seq (e.g. Drop-Seq). (4) Cells are lysed within droplets and the released mRNAs and SBOs are captured, reverse-transcribed, and sequenced. (5) Cells are demultiplexed and assigned to their origin and processed for further analysis. **(b)** Heatmap of normalized SBO counts for 6-plex human/mouse species-mixing experiment. Rows represent cells and columns represent SBOs. Cells are assessed whether they are positive for a particular SBO based on the SBO count matrix (see Online Methods). Cells were classified as singlets (positive for a unique SBO), multiplets (positive for more than one SBO), or negatives (not positive for any SBO) and ordered by their classifications. **(c)** Scatter plot showing raw counts between two SBOs. SBO 1 and 6 were used to barcode different samples (Human 1, Mouse 2) [left]. SBO 3 and 4 were used to barcode the same sample (Human 3) [right]. **(d)** Species-mixing plot of samples associated with SBO 1 and 5. Cells were labeled according to their SBO classification. Black dots indicate Human 1 sample barcoded with SBO 1, red dots indicate Mouse 1 sample barcoded with SBO 5, and grey dots indicate doublets that are positive for both SBOs. **(e)** Distribution of RNA transcript counts in cells between singlets (green), multiplets (blue), and negatives (red). Negatives which imply beads exposed to ambient RNA, had the lowest number of transcripts. Multiplets had slightly more transcripts than singlets, indicating more RNA content within a droplet.

To demonstrate our method’s ability and accuracy of multiplexing samples, we performed a 6-plex human/mouse species-mixing experiment. Two samples each of HEK293T cells and NIH3T3 cells carried a single unique SBO and one sample of each cell line carried a combination of the two SBOs **(Supplementary Fig. 2)**. We pooled all the samples together in equal proportions and performed a single run of Drop-Seq. Cells were deliberately overloaded during Drop-Seq to increase the chance of multiplets. We obtained 2,759 cell barcodes, in which at least 500 transcripts were detected and the cells were successfully assigned to their sample origins. Multiplets and negative cells were detected based on the SBO count matrix **(see Online Methods)**. Cells that were classified as singlets almost exclusively express their sample barcodes, while multiplets and negatives express multiple or no sample barcode, respectively (Fig. 1b). Scatter plots of SBO counts that originated from two different samples showed an exclusive relationship, whereas SBO counts from the same sample showed a strong correlation in their expressions (Fig. 1c, **Supplementary Fig. 3)**. Species classification using SBOs was consistent with the transcriptome-based species-mixing plot results (Fig. 1d, **Supplementary Fig. 4)**. We also observed a clear difference in the distribution of RNA transcripts between singlets, multiplets, and negatives as expected, indicating the unambiguous detection of multiplets and negatives (Fig. 1e). These results suggested that our method enabled sample multiplexing in single-cell experiments with high accuracy and specificity, and elimination of multiplets and negatives.

During drug discovery, gene expression profiling can be applied to annotate function of small molecules^7^ and to elucidate the mechanisms underlying a biological pathway^8, 9^. While systematic approaches to profile gene expression of a large number of small molecules using microarray technology has been made^10, 11^, they are limited to bulk measurements. To capture diverse responses of highly heterogeneous samples such as tumors, single-cell gene expression profiling is indispensable, although current technologies are not suitable for multiple screening. We envisioned that our method could be used when interested in screening multiple gene expression profiles in single-cells subjected to drug perturbations. We performed a 5-plex time-course scRNA-seq in the K562 cell line, which is derived from chronic myeloid leukemia and expresses the *Bcr-Abl* fusion gene^12^. We investigated the single-cell transcriptional response of K562 cells to imatinib, a BCR-ABL targeting drug^13^, over treatment time **(see Online Methods)**. Pseudotime analysis of single-cells in multiplexed samples collected from five time points showed a branched gene expression trajectory and a sequential progression in trajectory over drug treatment time (Fig. 2a). The branched trajectory showed that two transition states existed as a result of imatinib treatment. Samples exhibited asynchronous patterns in pseudotime, although the average increased with drug treatment time **(Supplementary Fig. 5c, d)**. Differential expression analysis over pseudotime identified several gene cohorts that change during the transition (Fig. 2b, **Supplementary Fig. 5a, b**). Notably, erythroid-related genes such as *HBZ* and *ALAS2* had increased in expression levels over pseudotime (Fig. 2b). This was consistent with previous studies that have shown increased expression of *HBZ* in imatinib-treated cells^14, 15^. Differentially expressed genes (DEGs) between the two transition states were also identified **(Supplementary Fig. 5e)**.

**Figure 2.**
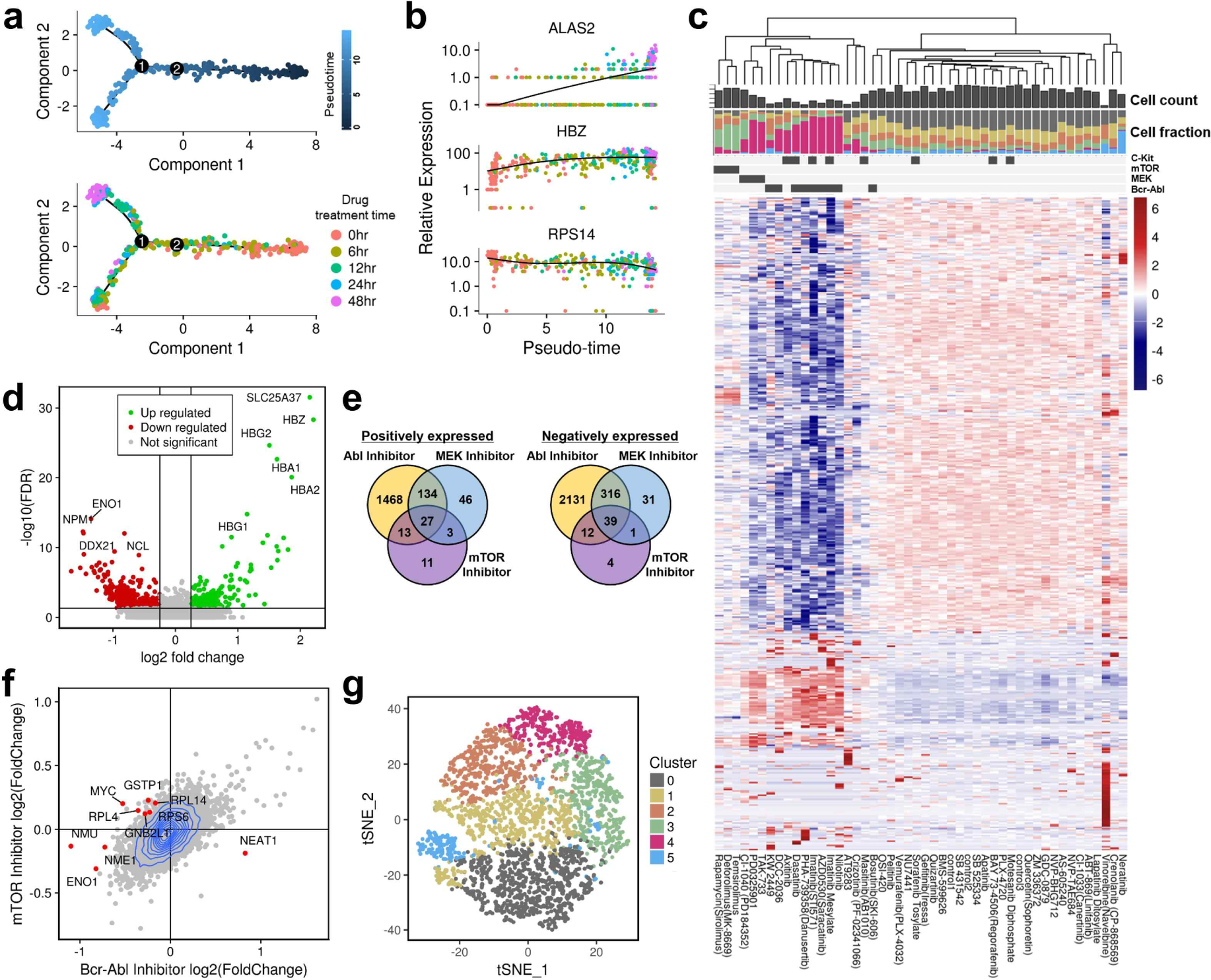
Applications of multiplexed single-cell RNA-seq using transient barcoding. **(a)** Monocle pseudotime trajectory of K562 cells treated with imatinib at different time points. Cells are labeled by pseudotime (top) and drug treatment time (bottom). **(b)** Prominent gene expression alterations in 5-plex time-course experiments of imatinib treatment. Note that the cells are labeled by drug treatment time and are not synchronously distributed over pseudotime. **(c)** Hierarchical clustered heatmap of averaged gene expression profiles for 48-plex drug treatment experiments in K562 cells. Each column represents averaged data in a drug and each row represents a gene. Differential expressed genes were used in this heatmap. The scale bar of relative expression is on the right side. The ability of the drugs to inhibit kinase proteins is shown as binary colors (dark grey indicating positive) at the top. The bar plot at the top shows the cell count for each drug and the bar plot at the bottom represents a relative fraction of cells in each t-distributed stochastic neighbor embedding (t-SNE) cluster (shown in Figure 2g). **(d)** Volcano plot displaying differential expressed genes of imatinib mesylate compared with DMSO controls. Genes that have a p-value smaller than 0.05 and an absolute value of log (fold change) larger than 0.25 are considered significant. Up-regulated genes are colored in green, down-regulated genes are colored in red, and insignificant genes are colored in grey. Ten genes with the lowest p-value are labeled. **(e)** Venn diagram showing the relationship between DEGs of three drug groups. Fourteen drugs are classified into three groups according to their protein targets (see Figure 2c, top) and differential expression analysis is performed by comparing each group with DMSO controls. Relations of both positively (left) and negatively (right) regulated genes in each group are shown. **(f)** Plot showing a correlation between fold changes of expression in cells treated with mTOR inhibitors and BCR-ABL inhibitors compared with DMSO controls. (g) t-SNE plot of a single-cell in the 48-plex K562 samples. Plot shows six clusters.

Next, we asked whether our approach could be used for simultaneous single-cell transcriptome profiling for multiple drugs in K562 cells. We selected 45 drugs, of which a majority was kinase inhibitors, including several BCR-ABL targeting drugs. Three dimethyl sulphoxide (DMSO) samples were used as controls **(Supplementary Table 1)**. A 48-plex single-cell experiment was performed by barcoding and pooling all samples after drug treatments. A total of 3,091 cells were obtained and demultiplexed after eliminating multiplets and negatives. The averaged expression profiles of each drug were visualized as a heatmap (Fig. 2c). Each drug exhibited its own expression pattern of responsive genes. Interestingly, unsupervised hierarchical clustering of the averaged expression data for each drug revealed that the response-inducing drugs clustered together by their protein targets, whereas drugs that induced no response showed similar expression patterns with DMSO controls (Fig. 2c, **Supplementary Fig. 6)**. Also, we can evaluate cell toxicity by examining the cell counts of each drug. Drugs that targeted BCR-ABL or ABL showed the strongest response and toxicity, and drugs that targeted MEK or mTOR showed relatively mild response. Differential expression analysis based on the single-cell gene expression data identified DEGs for each drug (Fig. 2d, **Supplementary Fig. 7, and Supplementary Table 3)**. We note that highly expressed erythroid-related genes such as *HBZ, HBA,* and *HBG* were up-regulated, and genes such as *DDX21, NCL, ENO1,* and *NPM1* were down-regulated in the sample treated with imatinib (Fig. 2d). Similar DEGs were identified for other drugs targeting BCR-ABL. Drugs such as vinorelbine and neratinib showed unique gene expression signatures and DEGs. Analysis of drugs grouped by their protein targets showed different relationships between DEGs of each protein target (Fig. 2e, **Supplementary Table 4**). In addition, comparative analysis of two groups revealed different expression profiles (Fig. 2f, **Supplementary Fig. 8**).

In order to comprehensively analyze the drug screening data at a single-cell resolution, we performed unsupervised clustering analysis on all the single-cell datasets. We observed six clusters (Fig. 2g), which were not clearly separated possibly due to a highly complex transcriptional space. Nevertheless, for each drug, the relative abundance of cells assigned to each cluster was various (Fig. 2c, **Supplementary Fig. 9c)**. Most of the cells affected by BCR-ABL and MEK inhibitors were concentrated in cluster 4, whereas cells affected by mTOR inhibitors were mainly concentrated in cluster 3. Especially, most of cells in cluster 5 belong to neratinib treated sample. Several markers associated with each cluster were verified by differential expression analysis **(Supplementary Fig. 10)**. Analysis of cell cycle states revealed no association between cell cycle states and specific clusters **(Supplementary Fig. 9a)**. The fraction of highly proliferative state (G2 phase) was decreased in samples treated with BCR-ABL targeting drugs possibly due to drug-induced cell cycle arrest^16^ **(Supplementary Fig. 9b)**. High-depth sequencing with more single-cells may detect more heterogeneous and distinct cellular responses for each drug.

To validate the universal applicability of our methods, we performed a 48-plex drug screening experiment on the A375 cell line (*BRAF* V600E-positive^17^) with an identical drug set. Similar to K562 cells, response-inducing drugs were clustered together in a target-specific manner in A375 cells **(Supplementary Fig. 11, 12, and Supplementary Table 5)**. Our results showed that multiplexed scRNA-seq could be used to screen single-cell transcriptional responses to drugs in a high-throughput manner and drug targets could be estimated by their transcriptional patterns.

We have developed a novel method for multiplexed scRNA-seq via transient transfection, which offers several advantages over currently available methods. Cost is substantially saved when performing scRNA-seq for multiple conditions. Batch effect, a major challenge in scRNA-seq^18^, can be significantly reduced by pooling and running all samples together, enabling more precise analysis between single-cell samples. Our method could eliminate multiplets and negatives, potentially able to increase throughput of scRNA-seq by using high concentration cells as an input. Removal of multiplets and negatives is also advantageous for single-cell analysis. Although a few methods for multiplexed scRNA-seq has been developed recently^19, 20^, which use natural genetic barcodes and antibody tagging, they have limited applications for genetically diverse samples^19^ and require valuable reagents and surface markers, respectively^20^. Our approach using SBO transfection (1) is very simple and readily applicable in individual labs, (2) has potential to be applied to nucleus samples, and (3) offers a high capacity for multiplexing by using different combinations of SBOs. We expect that our multiplexing strategy plays a central role in the widespread adoption of scRNA-seq including drug screening.

## Materials and methods

### Cell lines and cell culture

All cell lines were obtained from Korean Cell Line Bank (KCLB) and maintained at 37°C with 5% CO_2_. The human embryonic kidney HEK293T, the mouse embryo fibroblast MH3T3, and the human malignant melanoma A375 cell lines were cultured in Dulbecco’s modified Eagle’s medium (DMEM; Gibco, USA) supplemented with 10% fetal bovine serum (FBS; Gibco, USA) and 1% penicillin/streptomycin (Thermo Fisher Scientific, USA). The human chronic myelogenous leukemia K562 cell line was cultured in Roswell Park Memorial Institute (RPMI; Gibco, USA) medium supplemented with 10% FBS and 1% penicillin/streptomycin.

### Barcode design and transfection

The short barcoding oligo (SBO) contains a unique 8 bp sample barcode, an amplification handle, and a poly-A tail. 5’-TCCAAGGTACAGACCTCTGACGNNNNNNNN(A)_30_-3’ is the full SBO sequence. ‘TCCAAGGTACAGACCTATATCTGACG’ is the amplification handle sequence, ‘NNNNNNNN’ is the sample barcode sequence, and (A)30 is the poly-A tail sequence. All SBOs were prepared by IDT (Integrated DNA Technologies, USA) without any modifications. Four hours before Drop-Seq, 28 pmol/mL of SBO was transfected per well using Lipofectamine^™^ 3000 (Life Technologies, USA) according to the manufacturer’s protocol.

### Drop-seq: NGS preparation of mRNA and SBOs

For each experiment, samples of various conditions were pooled together. The pooled cells were passed through a 40-micron filter and diluted at a final combined concentration of 100~400 cells/μL according to Drop-Seq protocol instructions^1^. Droplets were generated and processed as previously described. Droplets were collected and the recovered beads were processed for immediate reverse transcription, followed by Exonuclease I treatment. The resulting cDNA was divided into appropriate number of tubes, amplified using KAPA HiFi HotStart PCR Kit (Kapa Biosystems, Inc., Switzerland). cDNA amplification was performed in 50ul PCR reaction which included 4 μL of 10 μM SMART PCR primer, 25 μL KAPA HiFi DNA polymerase, and up to 21 μL of nuclease-free water. Then, PCR reaction was performed using the following protocol: 3 min at 95°C; 4 cycles of (20 s at 98°C, 45 s at 65°C, 3 min s at 72°C); 9 cycles of (20 s at 98°C, 20 s at 67°C, 3 min s at 72°C); 5 min at 72°C. The PCR products were purified twice using 0.6X AMPure (Beckman coulter, USA) beads according to manufacturer’s instructions. To get reverse transcribed SBO that are much shorter than cDNA, the first supernatant from AMPure purification step was further purified adding 1.4X homemade AMPure beads (using Sera-Mag SpeedBeads (Thermo Scientific, USA), hereafter, Serapure beads^21^). The cDNA products were fragmented and further amplified using the Nextera XT DNA library preparation kit (Illumina, USA).

The SBO library preparation was performed using a two-step PCR protocol. One nanogram of the SBO cDNA product was loaded into 20 μL of the first adaptor PCR reaction, which included 1 μL of 10 μM forward and reverse primers, 10 μL KAPA HiFi DNA polymerase, and up to 8 μL of nuclease-free water. PCR reaction was performed using the following protocol: 3 min at 95°C; 8 cycles of (20 s at 95°C, 20 s at 64°C, 20 s at 72°C); 5 min at 72°C using the following primers: SMART+AC; P7-SBO hybrid. After 1.8X Serapure bead purification, 8 μL of the first PCR product was loaded into 20 μL of the second index PCR reaction, which included 1 μ of 10 μM forward and reverse primers, and 10 μL KAPA HiFi DNA polymerase. PCR reaction was performed using the following protocol: 3 min at 95°C; 6 cycles of (20 s at 95°C, 20 s at 60°C, 20 s at 72°C); 5 min at 72°C using the following primers: New-P5-SMART PCR hybrid; Nextera index Oligo. The second PCR product was purified using 1.2X Serapure beads. All primers were prepared by IDT. Sequencing was performed on the Illumina NextSeq 500 using NextSeq 500/550 High Output v2 kit (75 cycles) (Illumina, USA). The sequences of primers were provided in Supplementary Table 2.

### 6-plex human/mouse species-mixing experiment

HEK293T and NIH3T3 cells were prepared one day before Drop-Seq and plated on 6-well plates (TPP, Switzerland) at approximately 70% confluency. Transfection of 28 pmol/mL of SBO was performed 4 hours before Drop-Seq, as described above. All cell samples were trypsinized using Trypsin-EDTA (0.25%) and phenol red (Gibco, USA), pooled together, and washed four times with phosphate-buffered saline (PBS; Gibco, USA). The cells were then resuspended in 0.01% BSA + PBS, passed through a 40-micron filter, counted using LUNA^™^ Automated Cell Counter (Logos Biosystems, Korea), and diluted at a final combined concentration of 400 cells/μL. The diluted sample library was run once in Drop-Seq and sample preparation and sequencing were performed as above. From one Drop-Seq run, about 77,000 beads were obtained and divided into 24 PCR reactions for cDNA amplification. Sample preparation was completed using two reactions of the Nextera XT DNA library preparation kit (Illumina, USA).

### 5-plex time-course experiment of drug treatment

K562 cells were plated on 6-well plates at approximately 30% confluency. 1 μM imatinib was treated to K562 for five time points (0, 6, 12, 24, and 48 hours after treatment). Transfection of 28 pmol/mL of SBO was performed 4 hours before Drop-Seq, as described above and the cells of each condition were pooled, washed four times with PBS, and resuspended with 0.01% BSA + PBS. After filtering and counting, the pooled cells were diluted at a final combined concentration of 100 cells/μL. The diluted sample library was run once in Drop-Seq. Sample preparation and sequencing were performed as above. From one Drop-Seq run, the pooled beads were divided into 24 PCR reactions for cDNA amplification. Sample preparation was completed using two reactions of the Nextera XT DNA library preparation kit.

### 48-plex drug screening experiment in K562 cells

K562 cells were plated on 24-well plates at approximately 30% confluency and treated with 1 μM of each drug. After 44 hours, transfection of 28 pmol/mL of SBO was performed. Four hours after the transfection, the cell samples from each drug treatment were pooled. The diluted sample library was run three times in Drop-Seq. All subsequent steps were the same as described above in the 5-plex time-course experiment. After three Drop-Seq runs, the pooled beads were divided into 48 PCR reactions for cDNA amplification. Sample preparation was completed using three reactions of the Nextera XT DNA library preparation kit.

### 48-plex drug screening experiment in A375 cells

A375 cells were prepared one day before drug screening and plated on 24-well plates at approximately 30% confluency. All subsequent steps were the same as described above in the 48-plex drug treatment experiment. After three Drop-Seq runs, the pooled beads were divided into 48 PCR reactions for cDNA amplification. Sample preparation was completed using three reactions of the Nextera XT DNA library preparation kit.

### Single-cell transcriptome data processing

For each NextSeq sequencing run, raw sequencing data were converted to FASTQ files using bcl2fastq2 software (Illumina). Each sequencing sample was demultiplexed using Nextera N7xx indices. Tagging, trimming, alignment, and adding annotation tags were performed according to the standard Drop-Seq pipeline (http://mccarrollab.com/dropseq). Briefly, reads were first tagged according to the 12-bp cell barcode sequence and the 8-bp unique molecular identifier (UMI) in “read 1”. Then, reads in “read 2” were aligned with the hg19 or hg19-mm10 concatenated reference depending on the experiments and collapsed onto 12-bp cell barcodes that corresponded to individual beads. Hamming distance of 1 was used to collapse UMI within each transcript. Digital expression matrix was obtained by collapsing filtered and mapped reads for each gene by UMI sequence within each cell barcode.

### Sample barcode (SBOs) processing

FASTQ files of SBOs were generated as described above. Raw sequencing reads were trimmed to remove PCR handles. Cell barcodes and UMIs were extracted from read 1 and sample barcodes were extracted from read 2. Reads were assigned to 8-mer of sample barcode reference **(Supplementary Table 2)** with a single-base error tolerance (Hamming distance = 1) and Cell barcodes x sample barcodes count matrix (Hereinafter referred to as SBO matrix) was generated with consideration to UMI de-duplication. All the processes were made by our homemade python scripts.

### Merging Sample barcode and transcriptome data

Independently obtained cell barcodes from the two matrices (SBO matrix, transcriptome matrix) were compared and merged based on the cell barcodes from the transcriptome matrix. When merging, hamming distance of 1 was allowed. Finally, the columns of the SBO and the transcriptome matrix consisted of the same cell barcodes.

### Demultiplexing and classification of samples using SBO matrix

SBO matrix was normalized using a modified version of centered log ratio (CLR) transformation^22^:

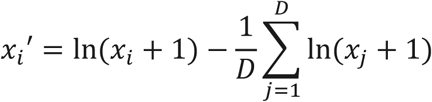

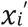 denotes the normalized count for a specific SBO in cell *i*, *x_i_* denotes the raw count, and *D* is the total cell number. In CLR transformation, the raw counts of SBO are divided by the geometric mean of individual SBO across cells and are log-transformed. We added the raw counts of SBO to 1 to avoid infinite values. We hypothesized that we can discriminate positive signals from negative (background) signals by fitting the distribution of negative signals of each SBO and thresholding the normalized counts to a specific value of each SBO. Following the normalization, for each SBO, we excluded the cells with the highest expression of the SBO among all SBOs. We fitted a negative binomial distribution to the remaining cells to obtain a distribution of negative signals. Next, we calculated a quantile with 0.99 probability to get the threshold value of each SBO. Cells that have higher SBO counts than the threshold value were considered as positives for that SBO. Cells were demultiplexed and classified into singlets, multiplets, and negatives based on the above results.

### 6-plex human-mouse species mixing experiment analysis

Transcriptome and SBO data processing were performed as described above. We obtained 2,759 of cell barcodes after filtering out cells with less than 500 transcripts. After the SBO matrix normalization and classification, we classified singlets as positive for one of SBOs, multiplets as positive for more than one SBOs, and negatives as positive for none of SBOs. For species-mixing plots in Figure 1d and Supplementary Figure 4, only singlets and doublets of the two specified SBOs were used.

### Pseudotime analysis

For pseudotime analysis of 5-plex time-course experiment, we applied the R package ‘Monocle 2’^23^. After removing multiplets and negatives, samples were demultiplexed, quality controlled, and analyzed. A single cell trajectory was constructed by Discriminative Dimensionality Reduction with Trees^24^ (DDRTree) algorithm using genes differentially expressed at different time points. Cells were ordered across the trajectory by setting the state containing 0 hour sample as a time zero and pseudotime was calculated. In order to identify differentially expressed genes (DEG) over pseudotime, a likelihood ratio test in the negative binomial model was performed and genes with a q-value less than 0.01 were selected as DEG. When drawing the heatmap, genes were clustered by their pseudotime expression patterns. Differential expression analysis between two transition states in branch 1 was performed using BEAM function in Monocle package.

### 48-plex drug screening data analysis

Following the alignment of sequencing reads, downstream analysis of the 48-plex drug screening experiment was performed using the R package ‘Seurat’^25^. After demultiplexing and removing multiplets and negatives, cells were quality-controlled based on the mitochondrial reads fraction, number of UMI, and number of genes. We identified 3091 cells in which at least 500 transcripts and 300 genes were detected. RNA expression matrix was log-normalized and processed for the further analysis. To cluster the single cells, we ran principal component analysis (PCA) using the expression matrix of variable genes, then performed t-distributed Stochastic Neighbor Embedding (t-SNE) using the first 6 PCA components. We identified six clusters using FindClusters function in Seurat with resolution=0.6. We assigned cell cycle phase scores using cell cycle markers^26^ and classified each cell to G2M, S, or G1 phase. To draw a hierarchical clustered heatmap, we first identified DEGs for each drug with adjusted p-value<0.05 by Wilcoxon rank-sum test and obtained 469 responsive genes by merging the DEGs altogether. Expression levels of each drug for the responsive genes were normalized, averaged, and scaled and were used for drawing the heatmap. To construct the dendrogram at the heatmap, hierarchical clustering was performed based on correlations among the expression levels across drugs. Normalized and scaled gene expression data were used in the heatmap. To identify DEGs in Figure 2d and Supplementary Figure 7, we performed likelihood ratio test between single cells in each drug and single cells in DMSO controls. DEGs with adjusted p-value<0.05 and |log 2 (fold change)|>0.25 were listed in Supplementary Table 3. To analyze samples by their protein targets, fourteen drugs were classified into three groups (BCR-ABL inhibitors, MEK inhibitors, mTOR inhibitors). Differential expression analysis between cells in each groups and cells in DMSO controls was performed as described above and DEGs were listed in Supplementary Table 4. The analysis of drug screening experiment of A375 was performed in the same manner as K562.

## Data availability

All data generated or analyzed during this study are included in this published article or will be provided by the corresponding authors upon request.

## Acknowledgements

This work was supported by the following sources: (i) the Mid-career Researcher Program (2015R1A2A1A10055972) through the National Research Foundation of Korea, funded by the Ministry of Science, ICT & Future Planning, (ii) the Bio & Medical Technology Development Program of the National Research Foundation (NRF) funded by the Korean government (MSIT; NRF-2016M3A9B6948494), (iii) the Bio & Medical Technology Development Program of the National Research Foundation (NRF) funded by the Korean government (MSIT; NRF-2018M3A9H3024850), and (iv) by the Ministry of Science, ICT and future Planning (grant number: NRF-2018R1A2B2001322). We thank Wan Namkung of the department of Pharmacy in Yonsei University for kindly contributing the kinase inhibitor library (Catalog No. L1200, Selleck Chemicals).

## Author contributions

D.S., W.L., J.H.L., and D.B. developed the concepts and designed the study. D.S. and W.L. performed the experiments. D.S. performed bioinformatic analysis and analyzed the data. D.S. and W.L. wrote the manuscript with feedback from all authors. J.H.L. and D.B. supervised the project.

